# Low light levels increase avoidance behaviour of diurnal fish species: Implications for road culverts

**DOI:** 10.1101/2020.01.07.896605

**Authors:** John K. Keep, Jabin R. Watson, Rebecca L. Cramp, Matthew J. Jones, Matthew A. Gordos, Patrick Ward, Craig E. Franklin

## Abstract

Inadequately designed culverts are known to pose hydraulic barriers to fish passage, but they may also be behavioural barriers if they adversely affect light levels within them. To test this, we performed a choice experiment and quantified the amount of time individuals of four Australian fish species spent in darkened and illuminated areas of an experimental swimming fume. Behavioural responses were reflective of the species’ diel activity patterns; diurnal species preferred illuminated regions, while nocturnal species preferred the darkened region. We then determined a threshold light level of only ~100-200 lux (c.f. midday sunlight ~100,000 lux) was required to overcome the behavioural barrier in ~ 70% of the diurnal fish tested. Placing these threshold values into field context, 100% of culverts sampled recorded inadequate light levels. Attention is required to better understand the impacts of low light levels in culverts on fish passage and to prioritise restoration.

## Introduction

Comprising less than one percent of surface waters, freshwater ecosystems support approximately half of all extant fish species (Reid *et al.* 2013). Yet, competition for, and misuse of, freshwater resources has led to a significant decline in fish diversity and abundance with approximately one third of assessed freshwater fish now at risk of extinction (Dudgeon *et al.* 2006; IUCN 2019). A loss of connectivity between freshwater environments (fragmentation) has been a significant contributor to freshwater fish declines (Baumgartner *et al.* 2014; Harris *et al.* 2017; Grill *et al*. 2019). Connectivity is intrinsically linked with access to resources (food, habitat), key population drivers (immigration, emigration, access to spawning grounds), predator avoidance (Harris *et al.* 2017; Rodgers *et al.* 2014; Watson *et al.* 2018), and increasingly with climate change, to find refuge pools during drought and to recolonise suitable habitat once flows return. A leading cause of freshwater habitat fragmentation is waterway infrastructure such as dams, weirs and road culverts (Grill et al. 2019). Traditionally, culverts were designed to move water underneath civil structures in an efficient and cost-effective manner, with little consideration given to the movement requirements of instream biota. Culverts can pose a physical barrier to fish movement by generating excessively high water velocities, excessive turbulence, and by creating a physical jump/drop through bed scouring (Goodrich *et al.* 2018; Rodgers *et al.* 2014; Watson *et al.* 2018). Additionally, culverts can act as behavioural barriers if conditions in and around the structure act to dissuade fish from passing through.

An emerging concern for fish passage is the potential for altered light levels (i.e. low light during the day or artificial light at night) in and around man-made structures to negatively influence fish movement and behaviour (Jones *et al.* 2017; Perkin *et al.* 2011). For most fish, vision is an important aspect of their sensory repertoire; visual systems are essential for orientation, breeding, foraging and predator avoidance. Fish behaviour is linked with diel light cycles, and it is increasingly apparent that anthropogenic disturbances to natural lighting regimes can have detrimental impacts on affected fish populations (Becker *et al.* 2013). Several studies have shown that artificial lighting at night (i.e. from street lights, transport networks, industry) can influence reproduction, community structure and movement in nocturnal fish (Becker *et al.* 2013; Riley *et al.* 2012; Ryer *et al.* 2009). Likewise, structures that limit natural light penetration can also alter the behaviour of diurnally active fish (Jones *et al.* 2017).

Although light is an important cue regulating the movement behaviour of many fish species, especially salmonids, the specific effects of light on fish passage are highly species-, life stage- and site-specific, and are often influenced by the presence of other behavioural or hydrodynamic stimuli (Banks 1969; Mueller and Simmons 2008; Vowles and Kemp 2012). Several reports have found that low light levels in covered structures (e.g. culverts, fishways, and weirs) can contribute to increased avoidance behaviour during the daytime downstream movements of salmon smolt (Kemp *et al.* 2006; Kemp *et al.* 2005; Kemp and Williams 2008; Tétard *et al.* 2019; Welton *et al.* 2002). Similarly, avoidance of darkened environments within covered fishways contributes to the reduced movement of several small-bodied Australian freshwater fish species (Jones *et al.* 2017). Abrupt changes in light intensity such as at the entrance/exits of fishways or culverts can also cause avoidance behaviour in lampreys (Moser and Mesa 2009) and juvenile salmon (Ono and Simenstad 2014). However, other studies have demonstrated that upstream migrating salmon, trout, eels, Topeka Shiner, Fathead Minnow and common galaxias in Australia, are unaffected by reduced light levels in civil structures (Fjeldstad *et al.* 2018; Amtstaetter *et al.* 2017; Kozarek *et al.* 2017; Gowans *et al.* 2003; Rogers and Cane 1979). These conflicting accounts of the effect of light on fish movement suggest a range of species-specific behavioural responses to different lighting conditions (Fjeldstad *et al.* 2018; Amtstaetter *et al.* 2017; Kozarek *et al.* 2017; Gowans *et al.* 2003; Rogers and Cane 1979), with such variability also indicating that our understanding of the effects of altered lighting regimes on fish movement is poor, despite this issue being raised in several fish passage guidelines (e.g. Fairfull and Witheridge 2003; Franklin *et al.* 2018).

Accordingly, research is required to better understand the potential for low light levels within culverts to impact fish movement and behaviour to inform the regulation of new culvert structures and to guide the remediation of existing structures. The aim of this study was to quantify the effect of reduced light levels on the movement behaviour of four species of small-bodied or juvenile Australian native fish. We chose two small-bodied species, Fly-specked Hardyhead *(Craterocephalus stercusmuscarum)* (Günther, 1867) and Australian Smelt (*Retropinna semoni)* (Weber, 1895), both of which have maximum adult sizes of 7 cm. We also included juveniles of two large-bodied species, Australian Bass *(Macquaria novemaculeata)* (Steindachner 1866) and Silver Perch (*Bidyanus bidyanus*) (Mitchell 1838) that have respective maximum adult sizes of 60 and 40 cm. Three of the species, Fly-specked Hardyhead, Australian Smelt and Silver Perch, are more active during the daytime (Baumgartner *et al.* 2008; Clunie and Koehn 2001; Mallen-Cooper 1999; Stuart and Mallen-Cooper 1999), while Australian Bass are generally crepuscular but can be active at other times of the day and night (Harris 1985; Smith *et al.* 2011). We hypothesised that Fly-specked Hardyhead, Australian Smelt and Silver Perch would prefer an illuminated environment, and that Australian Bass would prefer a darker environment. We then aimed to establish the minimum lighting thresholds for the species that displayed a preference for an illuminated environment. Finally, we placed these light threshold values into context by comparing them with light levels measured within existing culverts in south-east Queensland, Australia.

## Methods

### Fish collection and husbandry

Juvenile Australian Bass (*n* = 40; TL: mean ± SD 73.4 ± 8.7 mm;) and Silver Perch (*n* = 70; mean ± SD 65.95 ± 16.6 mm) were sourced from commercial hatcheries. Adult Fly-specked Hardyheads (*n* = 110; mean ± SD 48.9 ± 4.7 mm) were supplied by a commercial collector (Aquagreen, Howard Springs, Northern Territory), from the Howard River, Girraween Road Crossing, Northern Territory (12°31’51”S 131° 07’41”E). Adult Smelt (*n* = 60; mean ± SD 42.45 ± 6.5 mm) were collected using nets at Cedar Creek and Moggil Creek, Brisbane, Queensland (27°19’28.6”S 152°47’39.1”E and 27°30’16.1”S 152°55’50.1”E, respectively).

The fish were housed at the Biohydrodynamics Laboratory at the University of Queensland (Brisbane, Queensland, Australia). Fish were kept with conspecifics in 40 L glass aquaria that formed part of a 1000 L recirculating system with mechanical and biological filtration and UV sterilization. The water temperature was maintained at 25°C ± 1°C. Fish were fed commercial aquaculture pellets (Ridley, Brisbane Australia) and exposed to a 12-hour light-dark cycle provided by overhead LED aquarium lighting. The ambient light intensity was measured at the water level of the housing aquaria using a photometer (Extech HD450, New Hampshire, U.S.A.), which averaged 2535 ± 238.6 lux (mean ± SD).

### Light – dark behavioural trials

Behavioural trials were performed on all four species in a 12-metre hydraulic channel (12.0 × 0.5 × 0.3 m). The light around and within the channel was controlled using blackout plastic sheeting to create an environment with zero ambient light (0 lux). The integrity of this screen was checked before starting trials each day to ensure no external light sources were present. Half of the channel was illuminated using 4000 K correlated colour temperature LED lighting (Atom 56-watt batten, China) and the other half left darkened. The light intensity above the illuminated half was set to 2535 lux, the same as above the housing aquaria. A sharp light-dark transition point was achieved by dividing the darkened area around the channel with black plastic. This included the space within the channel above the waterline.

Four treatments were required to control for the direction of water flow in the channel that could not be changed, and the illuminated state (light or dark) of the release point (Fig. 1). The first two treatments were with the downstream half of the channel illuminated and the upstream dark. Ten fish per species were randomly allocated to each treatment for each trial. The time each fish spent in each zone of the channel was observed for 30 min through a small observation point at the transition zone between the illuminated and darkened areas. All fish were swum individually and were released 1 m from either end of the channel, facing into the water flow (Fig. 1). The channel bulk velocity was set to 0.3 m s-1 and the depth set to 0.15 m, measured at the mid-point, 6 m along the channel length. This velocity was chosen as it was significantly below the maximum sustainable swimming speed (*U*crit; Brett 1964) of all four species tested (Watson *et al.* 2019) so as to minimise any effect that swimming capacity, or their innate rheotactic response, could have on their subsequent behaviour. The water temperature was maintained at 25 ± 1°C.

**Figure 1.**
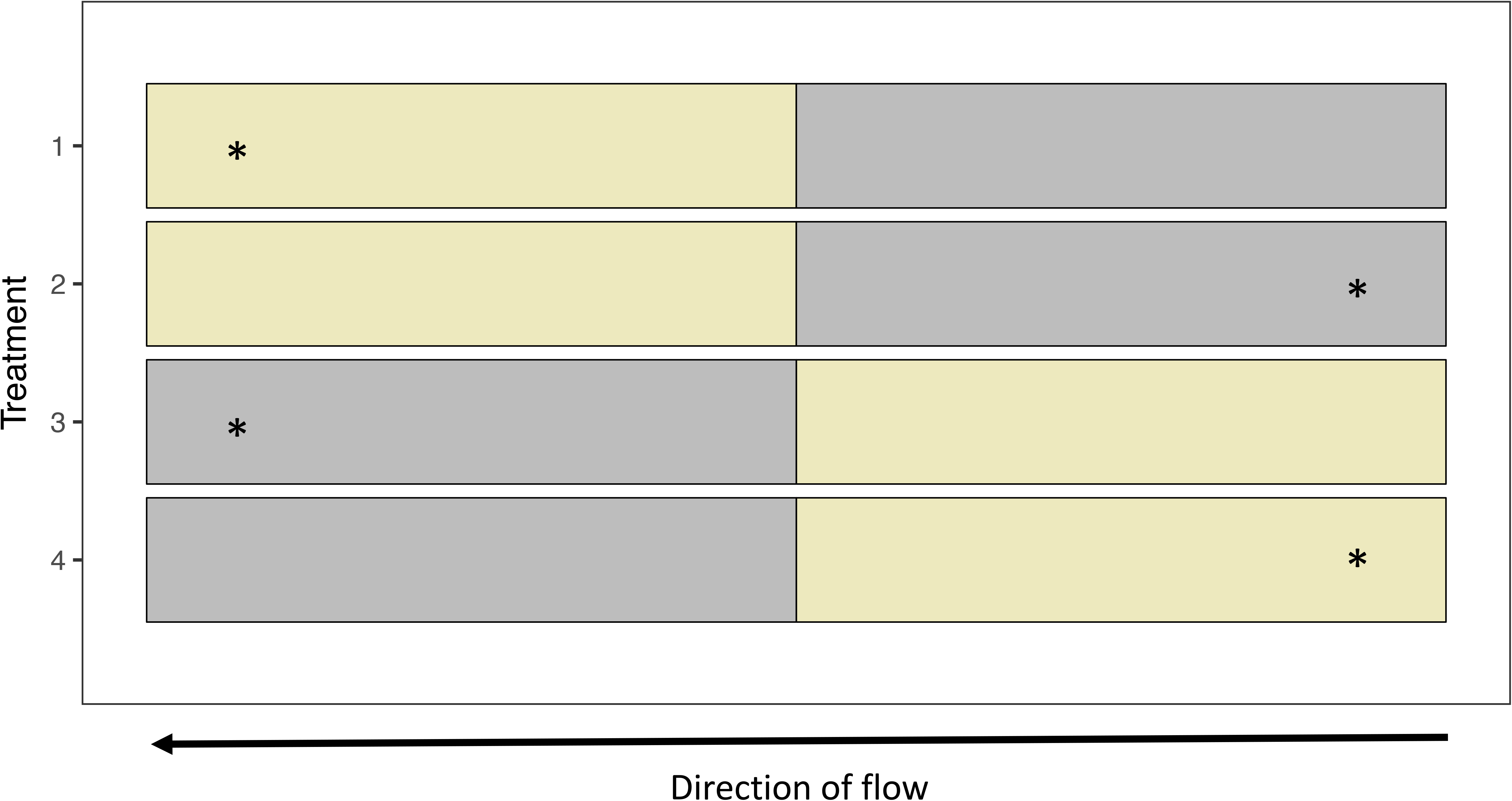
Schematic of experimental design to assess light preferences. Side view of the 12 m long experimental hydraulic flume, indicating the position of release (asterisks) and location of the darkened (grey) and illuminated (yellow) zones in relation to the direction of water flow.

### Determining the light intensity thresholds for fly-specked hardyheads and smelt

Of the four species tested, the Fly-specked Hardyhead and Australian Smelt displayed strong avoidance of darkened environments which could negatively impact their movement through artificially darkened culverts. To understand the minimum illumination levels that would encourage Fly-specked Hardyhead and Australian Smelt to enter a darkened environment, we set up the flume with the downstream half illuminated and upstream half darkened, and sequentially increased the light levels in the darkened section of the flume. The illuminated zone was set to the same light intensity as the housing aquaria (2535 lux). The darkened half of the channel was fitted with an overhead controllable LED strip light (ML-1009FAWi, MELEC, Birtinya, Queensland, Australia) that allowed us to incrementally increase light intensities in the darkened region. The light intensity treatments were determined by the response times as the experiment progressed. Fly-specked Hardyhead were released at 5, 10, 25, 50, 250, 300 and 400 lux, and Australian Smelt were released at 2.5, 5, 25 and 200 lux. We recorded the total time fish spent in both halves of the channel over 10 min, and the proportion (as a percentage) of individuals that used the darkened region of the flume (for any length of time). Fifteen fish of each species were individually tested in each light intensity treatment and released 1 m from the downstream end of the flume. Individual fish were only swum once. This was not done for Australian Bass which prefered the dark, or Silver Perch that showed no light-dark preference.

### Sampling light levels of culverts

To place the light intensity threshold values obtained for Fly-specked Hardyhead and Australian Smelt into context, we sampled the ambient light levels within and outside fifteen culverts within south-east Queensland (Australia) using a photometer (Extech HD450, New Hampshire, U.S.A.). Sampling was undertaken between 09:00 and 12:15 on the 16 December 2018 (austral summer), on a cloudless day when ambient light levels within the culvert would be at, or close to, their maximum levels. The culverts sampled were predominantly dual carriage roadways and one single pedestrian crossing (culvert range 3.4 – 7.0 m in length, ~ 1.0 m height). All culverts contained at least 0.2 m water depth at the time of sampling.

### Data analyses

All statistical analyses were performed using R version 1.1.423 (R Core Team 2017) in the RStudio environment. The preference experiment data was analysed using a quasibinomial generalised linear model and ANOVA with species, release condition (light or dark) and release point (downstream or upstream) as predictors, allowing for possible interactions. To analyse how many fish were entering the treatment zone, a binomial linear regression was fit to the data with ‘entering’ (y/n) as the response variable, and the time of day the fish were swum, species, light level in the treatment zone (lux) and fish length as predictors allowing all interactions. A subsequent ANOVA revealed that the light level in the treatment zone was the only significant predictor and the model was reduced accordingly. Statistical significance for all analysis was set at *P* < 0.05.

## Results

### Light – dark preferences

There was a statistically significant 3-way interaction between species, release point and lighting condition of the release point (F_3, 144_ = 11.2284, *p* < 0.001) due to the behaviour of Silver Perch (Fig. 2). When released downstream in the darkened environment, Silver Perch swam upstream into the illuminated zone, and when released downstream in the light they spent they spent around half of their time in the light. When released upstream they were indifferent in their lighting preference and stayed in the upstream zone.

**Figure 2.**
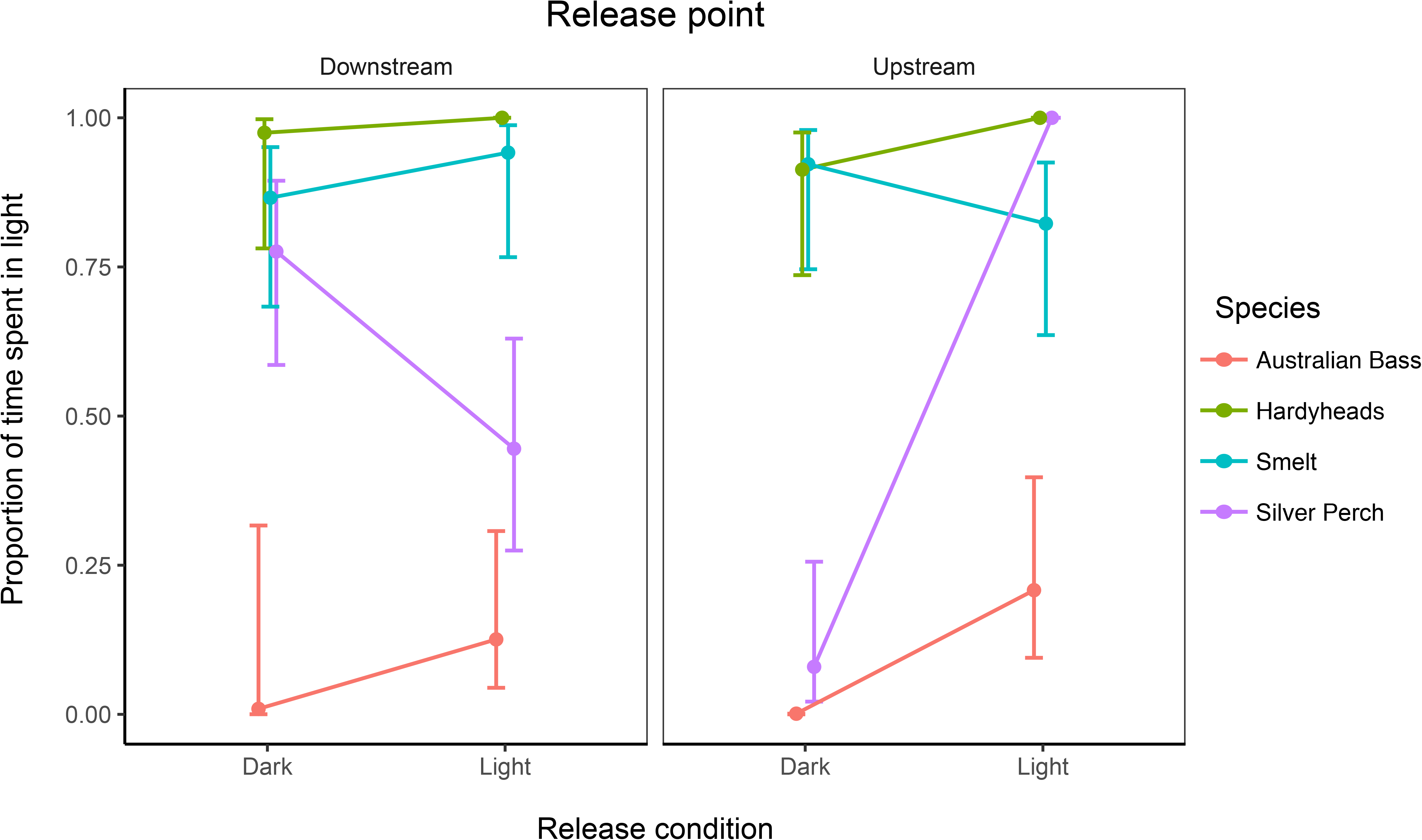
The average time spent in the light half of the experimental channel for each species and treatment, showing the statistically significant 3-way interaction was due to the behaviour of the Silver perch. Fly-specked hardyheads and smelt showed a strong preference for the illuminated region of the channel regardless of release point or its lighting conditions. Conversely Australian bass strongly preferred a dark environment regardless of release point or its lighting condition. Error bars represent 95% confidence intervals.

Australian Bass appeared to prefer the darkened zone of the channel, spending on average 91.3% of their time there across all treatments. Irrespective of flow direction, individual Australian Bass that were released in the illuminated zone (treatments 2 and 3) rapidly moved to the darkened zone. Neither the release point, nor the illumination condition at the release point were found to affect the time spent in either the light or dark zones.

In contrast, the Fly-specked Hardyhead and Australian Smelt both displayed a strong avoidance of the darkened zone (or preference for the light). Fly-specked Hardyhead and Australian Smelt were observed spending respectively 97.2% and 86.3% of their total trial time across all treatments in the illuminated zone. Like Australian Bass, neither the release point, nor the illumination condition at the release point were found to affect the time spent in either the light or dark zones. We observed that both the Fly-specked Hardyhead and Australian Smelt quickly moved to the illuminated zone of the channel when released in the darkened zone.

### Light intensity thresholds stimulating fish movement

Given that both Fly-specked Hardyhead and Australian Smelt displayed strong avoidance of the darkened environment in the channel, we gradually increased the illumination in the darkened (treatment) zone to determine the light threshold that would encourage these species to enter. Overall, the number of individuals entering the treatment region of the channel increased with increasing illumination (F_(1, 168)_ = 28.921, *p* < 0.001) (Fig. 3). It is worth nothing that while fish length did not have a statistically significant effect on the number of individuals entering the darkened treatment zones, length is potentially biologically significant with more larger fish entering at lower light levels (*p* = 0.056).

**Figure 3.**
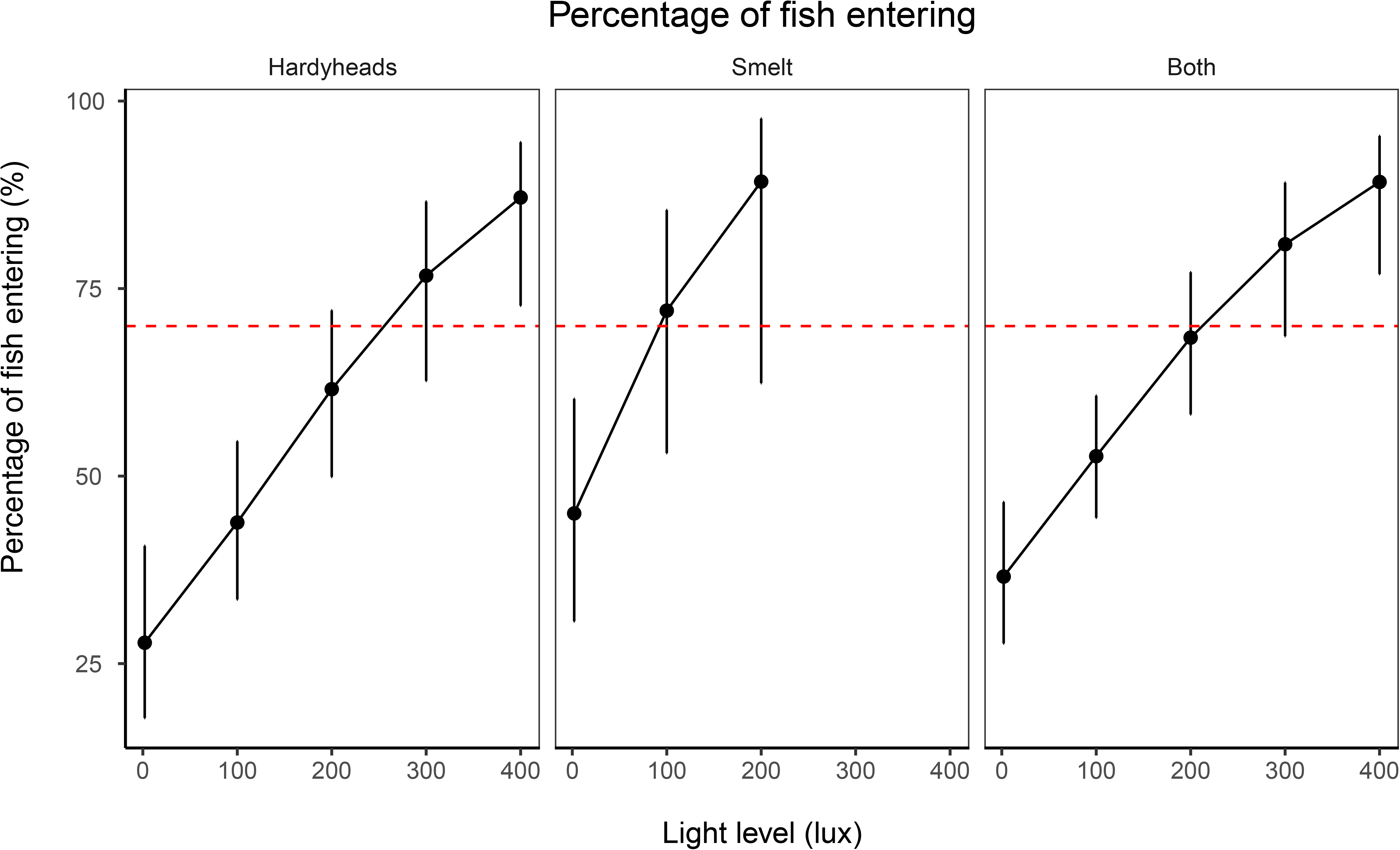
Regression curves showing the probability of a fish entering the darkened half of the experimental channel with increasing light levels. Both species were modelled individually and combined. Smelt were only tested up to 200 lux. Error bars represent 95% confidence intervals.

Fly-specked Hardyhead began entering the darkened region of the flume at 5 lux, with 26% of individuals observed entering. The number of individuals entering the darkened region of the flume remained at less than 50% until illumination levels exceeded 200 lux. The illumination threshold at which smelt started to enter the darkened region was 2.5 lux (13% of individuals). Doubling the amount of available light from 2.5 to 5 lux resulted in a four-fold increase to 53% of individuals entering. Further increasing the light intensity to 25 lux resulted in more than 75% of individuals entering the treatment zone.

### Light intensity responses in field context

We quantified the illumination levels in 15 culverts in south-east Queensland to determine how many reached the minimum lighting thresholds required to encourage 70% of Australian Smelt and Fly-specked Hardyhead to successfully move into a darkened environment. The modelled threshold values corresponded to 100 and 200 lux for Australian Smelt and Fly-specked Hardyhead, respectively. We found that lighting levels at the culvert entrance/exit averaged 70.9 ± 44.8 lux (mean ± s.d.; range: 5.6 - 123.1 lux; Table 1). In all culverts, light levels dropped to less than 3 lux in the centre (0.6 ± 0.8 lux; range 0 – 2.3 lux). Based on the light threshold determined for both Australian Smelt and Fly-specked Hardyhead, all culverts sampled contained insufficient light in the centre to promote a 70% passage success rate.

**Table 1:**
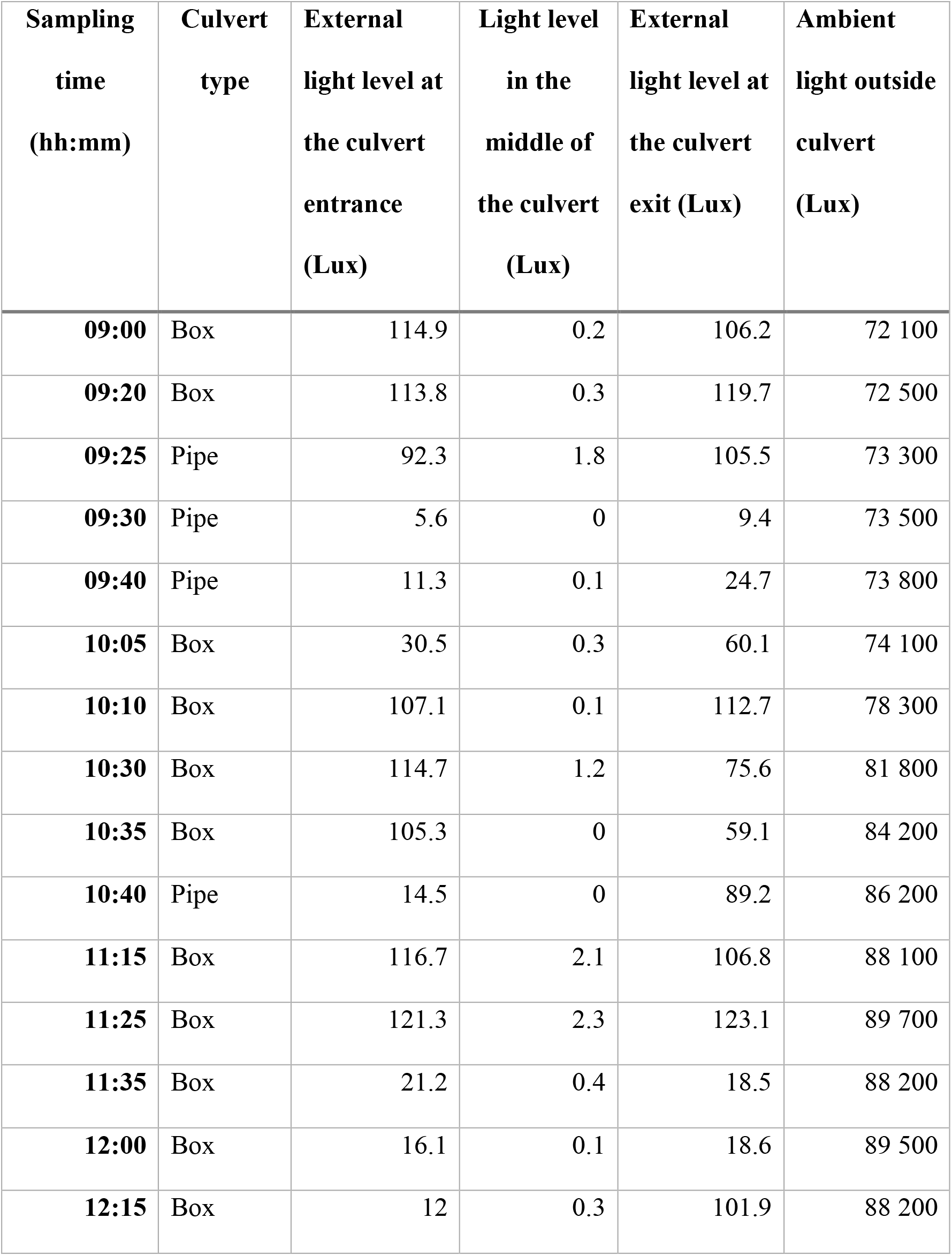
Light readings recorded at 15 roadway culverts.

## Discussion

Here we show that the levels of available light significantly affected the behaviour of three out of the four Australian fish species examined, and that our results were mostly consistent with the hypothesis that species behavioural responses to lighting levels would relate to their daily activity patterns (i.e. diurnal versus nocturnal). The largely diurnal Australian Smelt and Fly-specked Hardyhead showed near absolute avoidance of the completely darkened environment within the experimental channel, while the nocturnal Australian Bass strongly avoided the illuminated section. Surprisingly, Silver Perch showed no preference for either the illuminated or darkened environment. Silver Perch activity patterns in the wild are generally greatest during daylight hours, however, they do not appear to be actively inhibited by darkness and can be trapped, albeit at lower frequencies, at night (Baumgartner et al. 2008). In the present study, we were unable to disentangle a behavioural response to light levels from their response to water flow direction (rheotaxis), which may have been exacerbated by the relatively slow flow velocities used in this study compared to their swimming capacity (Watson *et al*. 2019). For the two species that avoided the darkened environment, we found that the threshold light intensities needed to encourage individual fish to enter the darker half of the test channel were quite low. These data also suggest that providing even very low levels of light with artificial lighting could remove the behavioural barrier that culverts may pose to diurnally active fish species.

The behavioural response of Australian Smelt, Fly-specked Hardyhead and Australian Bass provided an indication of the range of responses of fish species to low light levels representative of those within culverts, and how broad diel classifications can help to predict behavioural responses of those species most at risk of low light levels. Yet, consideration must be given to factors other than diel classification, such as movement motivation. Diadromous species that are obligate migrators for example, may be more likely to pass through a darkened culvert due a fundamental requirement to reach the sea or freshwater, as compared with facultative migrators. Indeed, the movement behaviour of *Galaxias spp*. was unaffected by a 70 m long darkened (0 lux) pipe culvert along an upstream migration path (Amtstaetter *et al.* 2017). In contrast, facultative migrators such as smelt, may lack that motivation and are more susceptible to altered light regimes (Jones *et al*. 2017). Clearly fish species differ in their readiness to use low light environments and so appropriate consideration of interspecific differences and variation in their tolerance of darkness within man-made structures needs to be given.

Abrupt lighting changes at sharp transitional point (as opposed to a graded transition) can be why some fish display strong behavioural reactions to light levels in some fish passage structures. Some fish may avoid areas where shadows cast by anthropogenic structures cause abrupt lighting changes because of the risk of predators using the shaded areas as cover (Kemp *et al.* 2005; Ono and Simenstad 2014; Steenbergen *et al.* 2011). However, data from our study, which also employed a sharp transition from light to dark, suggests this may not have been a constraining factor influencing the movement of the four fish species we examined. We found that amongst the species that showed a distinct light-dark preference response, nearly all individuals rapidly moved to their preferred illumination zone, regardless of the flow orientation or illumination state at the release point. While the sharp light gradient did not appear to completely restrict their initial movement into or out of the dark zone, further work will be required to determine if the abrupt light-dark transition influenced subsequent use of the space by the fish.

To determine the prevalence of prohibitively low light levels for fish passage in culverts, we measured light levels in 15 box or pipe culverts in Brisbane, Australia ranging from 3-8 m in length. We found that light levels at both the entrance and exit of the culvert ranged from ~5 to ~120 lux. Less than 3 lux was recorded in the middle of all culverts irrespective of culvert length. Based on the lighting thresholds for Fly-specked Hardyhead and Australian Smelt, all of the culverts examined could pose as a behavioural barrier to these species. Although we only measured light levels on just one day at each culvert, we measured at the brightest time of day (morning) and year (summer), so if prohibitively low light levels were detected under these conditions, they are likely to be light barriers at other times of year. The amount of ambient light present within the structure is dependent upon a culverts’ cross-sectional area, height and orientation relative to a light source. Environmental factors such as season, water depth and surrounding riparian vegetation density, will also affect the amount of light within a culvert, which means that the level of ambient light in a structure may vary considerably over daily and seasonal scales. A greater understanding of how culvert lighting conditions change over time is important for determining if and when a particular structure is likely to be a behavioural movement barrier for fish. Culvert lighting requirements need to be considered in the context of other culvert design features to ensure that efforts to mitigate hydrological barriers to fish passage (e.g. significant slope or excessive water velocities) do not inadvertently create behavioural obstacles to fish passage.

When assessing if lighting levels are likely to influence fish passage through culverts, it is important to consider other factors that may influence fish behaviour such as the presence of predators and food (Magurran 1990; Morgan and Godin 1985), schooling effects with conspecifics (Krause *et al.* 2000) or individuals’ personality (Hirsch *et al.* 2017). Our study was conducted under controlled laboratory conditions focusing on the test species’ response to light levels. Progressively overlaying the levels of complexity found in the field would strengthen our understanding of when reduced light levels are barriers, and how they may be overcome. For example, increases in water flow may reach a threshold point where a low light level ceases to be a behavioural barrier simply due to a strengthening of the fishes rheotactic response. Finally, light pollution at night from anthropogenic sources (Holker *et al.* 2010; Perkin *et al.* 2014) should be considered as increased levels of artificial ambient light along waterways may influence the movement behaviour of nocturnal species.

Currently, many fish passage guidelines for road crossing structures identify low light as a potential barrier for fish passage and recommend that light levels be considered by infrastructure planners and asset owners (Fairfull and Witheridge 2003; Franklin et al. 2018). Where light levels in culverts are predicted to impede passage of target fish species, alternate road crossing structures (such as bridges) may be recommended. However, in circumstances where low daytime light levels within a culvert may be unavoidable, our data suggest that providing small amounts of light though the installation of artificial lights or the provision of skylights, could remove a behavioural obstacle for some diurnally active fish species. More data on lighting thresholds for the movement of a greater range of fish species will inform fish passage guidelines and allow recommendations to be made in a site- and species-specific manner dictated by the culvert length, orientation, and the passage requirements of the local fish community.

## Conclusions

Our study showed that light levels affected the movement behaviour of three Australian native fish species, and that low light levels impeded the movement of two of the four species in an experimental channel. Optimal lighting levels for fish passage should be considered in the future design of artificial instream structures such as culverts and fishways, and in the remediation of existing structures. Our results indicate that only relatively low ambient daytime light levels are required within closed structures to encourage movement by certain diurnal species, and that fish willingly move into a darkened environment with the provision of artificial light. Developing minimum lighting standards that take into account species-specific light requirements can lead to improved passage rates through culverts, reduced fragmentation, and more resilient fish populations.

## Acknowledgements

The Research was funded by a National Environmental Science Program grant from the Threatened Species Recovery Hub to CEF [3.3.7] and an Australian Research Council Linkage Grant to CEF and MG [LP140100225]. This work was conducted with the approval of the University of Queensland Ethics Committee (SBS/312/15/ARC). We declare no conflicts of interest.

